# Early postpartum development of pup urine preference in mothers

**DOI:** 10.1101/2025.09.01.673527

**Authors:** Valentine Andreu, Rajyashree Sen, Nour El Houda Mimouni, Eun Ji Lee, Dianne-Lee Ferguson, Alexis Stutzman, Bianca J. Marlin

## Abstract

The transition to motherhood involves profound physiological and neural changes, including adaptations in the sensory systems that support infant care.^1,2^ While the olfactory system plays a critical role in guiding maternal behaviors such as pup retrieval and nesting,^3,4^ how olfactory processing itself is reshaped during motherhood remains poorly understood. Here, we show that first-time mothers develop a selective preference for pup urine following parturition and early postpartum care, a preference not observed for other social or neutral odors. Using odor preference assays combined with liquid and gas chromatography–mass spectrometry, we identify specific volatile compounds in pup urine that may contribute to this maternal attraction. Disruption of olfactory input or restriction of contact chemosensation abolished the preference, indicating that both volatile and non-volatile sensory modalities contribute, likely through combined input from the main olfactory epithelium (MOE) and vomeronasal organ (VNO).^5,6^ Notably, this preference is absent in late-pregnant females, in mothers separated from pups at birth, and in virgins cohoused with pups or exposed to pup urine-highlighting that pup urine preference depends on the convergence of internal hormonal signals and external chemosensory cues.^7,8^ These findings reveal a previously unrecognized specificity in maternal olfactory behavior and provide insight into how motherhood modulates the sense of smell to support offspring recognition and care.

**In Brief:** Maternal pup urine preference depends on both hormonal changes and chemosensory cues associated with motherhood.

**Highlights:**

- Pup urine specifically attracts postpartum mothers but not virgin females
- Pup urine contains distinct volatile and non-volatile metabolites
- Pup urine preference requires both pup experience and the hormonal priming of motherhood

## Results

### Pup urine specifically attracts postpartum mothers but not virgin females

Pup odors, specifically pup urine, are thought to guide maternal behaviors, such as licking and nest building.^9,4^ To investigate maternal behavior guided by pup odors, we conducted an approach-avoidance assay using urine from 5-day-old pups as the olfactory stimulus. We tested first-time postpartum day 5 (PPD5) mothers, who robustly exhibit maternal behaviors, and virgin females. All animals were first habituated to a tri-chamber apparatus. Twenty-four hours later, they were presented with two cotton swabs: one soaked in unfamiliar pup urine and the other clean (control), placed on opposite sides of the chamber (Figure 1A). To quantify interactions with cotton stimuli, we tracked the nose, tail-base, and center points of the cotton stimuli using DeepLabCut, a pose estimation software (Figure 1B–C).

**Figure 1.**
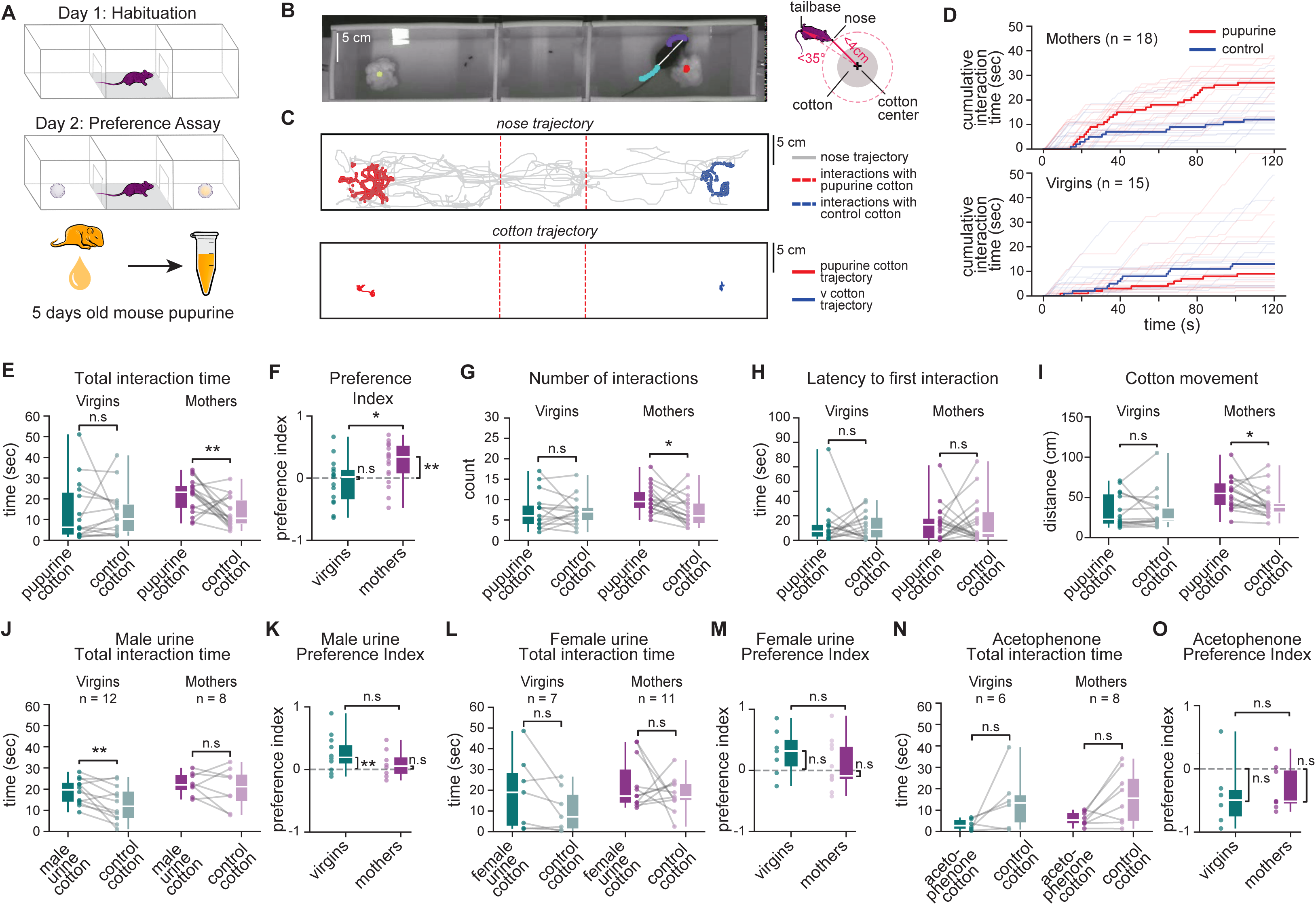
**PPD5 mothers selectively prefer pup urine.** (A) Odor preference tri-chamber assay set-up. (B) Left: Example camera frame from a representative session. Right: Schematic showing distance and angle thresholds used to define nose-directed interactions with each cotton. (C) Top: Representative nose trajectory (gray) from a single 2-min session. Colored dots mark keypoints tracked using DeepLabCut. Red and blue dots indicate interactions with pup urine-soaked and blank cotton, respectively. Bottom: Corresponding trajectories of the pup urine and blank cotton. (D) Cumulative interaction time with pup urine (red) vs. blank cotton (blue) over 2 min in PPD5 mothers (top; *n* = 18) and virgin females (bottom; *n =* 15). Bold lines represent the median across animals at each frame; fine lines show individual animals. (E) Total interaction time with pup urine vs. blank cotton in PPD5 mothers (n = 18) and virgins (n = 15). Box-and-whisker plots show minimum, 25th, 50th (median), 75th percentiles, and maximum. Distributions were compared using two-sided Wilcoxon signed-rank tests. In all panels, *p* < 0.05; *p* < 0.01; *n.s.*, not significant (*p* > 0.05). (F) Preference index (see Methods) calculated from data in (E). Distributions were compared against zero using the Wilcoxon signed-rank test (p-values shown on the side). Comparisons between cohorts were made using the Mann–Whitney U test (p-values shown above the plots). (G) Number of interactions for pup urine vs. blank cotton in PPD5 mothers (*n* = 18) and virgin females (*n* = 15). Distributions were compared using two-sided Wilcoxon signed-rank tests. (H) Latency to first interaction for pup urine vs. blank cotton in PPD5 mothers (*n* = 18) and virgin females (*n* = 15). Distributions were compared using one-sided Wilcoxon signed-rank test to assess whether test mice interacted significantly faster with pup urine-soaked cotton compared to blank cotton. (I) Total distance the pup urine and blank cotton were moved by PPD5 mothers (*n* = 18) and virgin females (*n* = 15). Distributions were compared using two-sided Wilcoxon signed-rank tests. (J-O) Interaction time and preference index for additional social and non-social odors: male urine (J-K; PPD5 mothers *n* = 8, virgins *n* = 12), female urine (L-M; PPD5 mothers *n* = 11, virgins *n* = 7), and acetophenone (N-O; PPD5 mothers *n* = 8, virgins *n* = 6). For all preference index panels in these and later figures, distributions were compared against zero using the Wilcoxon signed-rank test (p-values shown on the side). Comparisons between cohorts were made using the Mann–Whitney U test (p-values shown above the plots). For all pairwise comparisons of latency to first interaction in later figures, a one-sided Wilcoxon signed-rank test was used to assess whether test mice interacted significantly faster with stimulus cotton compared to blank cotton. For all pairwise comparisons of total interaction time, number of interactions, and cotton movement distance, two-sided Wilcoxon signed-rank tests were used.

PPD5 mothers showed a significantly stronger preference for the pup urine-soaked cotton compared to virgin females (Figure 1D–G & Video S1). To quantify this preference further, we computed a preference index, calculated as the difference in time spent with the pup urine and control cotton, divided by the total time spent with both. A positive score reflects a preference for pup urine, while a score of zero indicates no bias. Mothers exhibited significantly higher preference index values than virgins (Figure 1F), indicating a robust attraction to pup urine.

Consistent with this, mothers spent more time inspecting the pup urine cotton (Figure 1D–E), interacted with it more frequently (Figure 1G), and moved it more than the clean cotton (Figure 1I). In contrast, virgin females showed no significant preference in time spent, number of interactions, or cotton movement (Figures D-G, I). Neither group differed in latency to first interaction with either cotton sample (Figure 1H). During the habituation phase, when no olfactory stimuli were present, neither group showed a side preference. These findings suggest that the preference for pup urine is acquired during the transition to motherhood, rather than being innate.

To determine whether the maternal preference extended to other social odors, we tested both virgin females and PPD5 mothers for their responses to adult male and female urine. Male urine has previously been shown to be attractive to virgin females, and as expected, virgins displayed a preference for it (Figure 1L-M, Figure S1A). Mothers, however, exhibited a distinct pattern: although they interacted more frequently with the male urine-treated cotton (Figure SA2), the interaction bouts within a session were shorter, resulting in no overall time preference and a preference index that was not significantly different from zero (Figure 1J–K). Both virgins and mothers showed low latency to engage with the male urine, (Figures S1A4), suggesting that the volatiles in male urine may diffuse more rapidly and broadly than those in pup urine under our assay conditions. Notably, although virgin females showed a significant within-group preference for male urine and mothers did not, there was no significant difference in preference index between virgins and mothers, indicating that this metric may not fully capture nuanced changes in interaction dynamics across groups (also see Figure 2F & 2K). Further, neither PPD5 mothers nor virgin females showed a specific preference for female urine in time spent, number of interactions, latency to first interaction or cotton movement (Figure L-M, Figure S1B).

**Figure 2.**
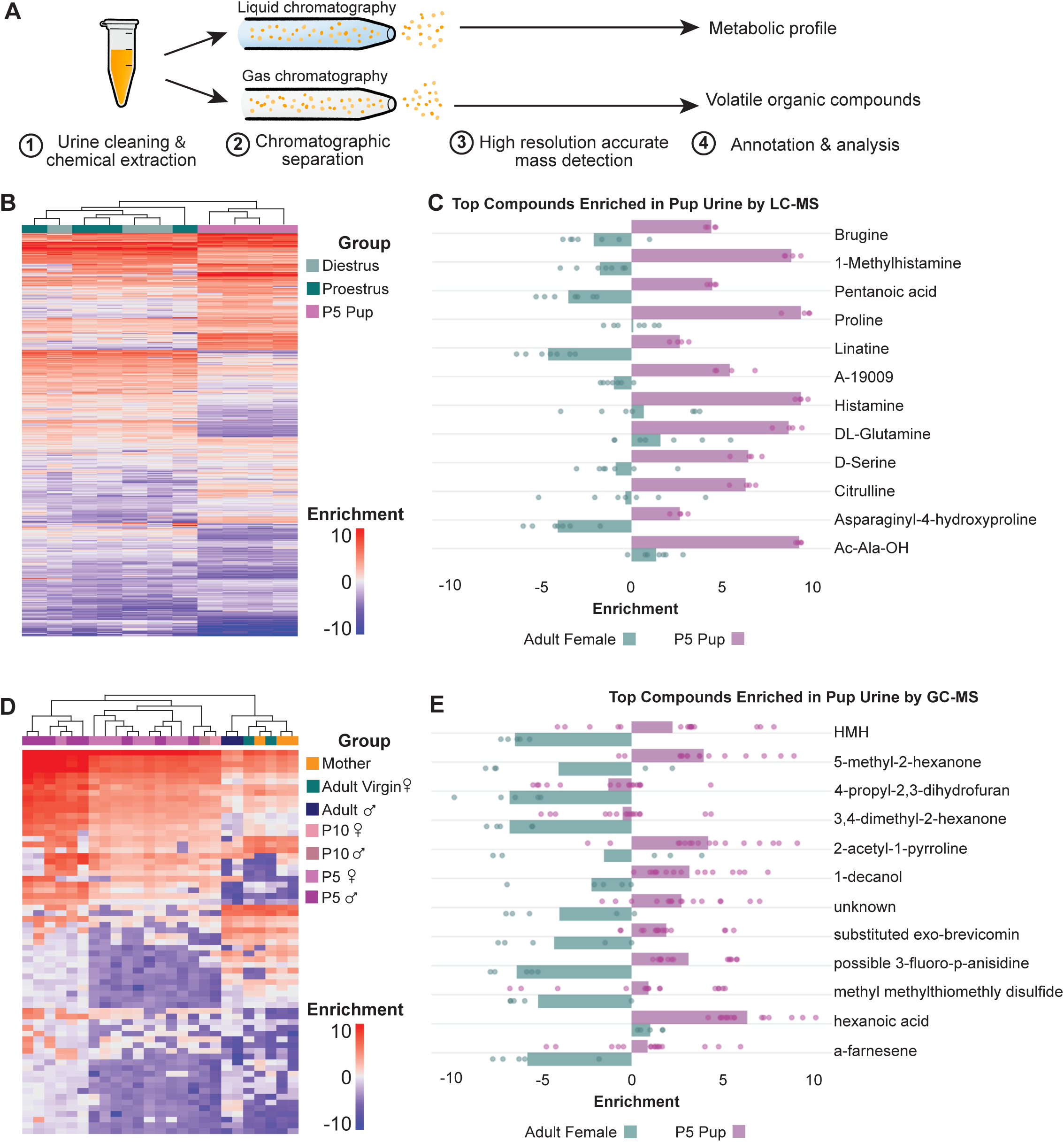
**Organic compound and metabolite signatures differ in adult and pup urine.** (A) Workflow and comparison of pup and adult female urine analyzed using liquid chromatography–mass spectrometry (LC-MS) and gas chromatography–mass spectrometry (GC-MS). Urine was pooled from 5-day-old pups (mixed-sex, same litter; *n* = 4) and 6–10-week-old virgin females in diestrus and proestrus (*n* = 4). (B) Supervised clustering of the all compounds analyzed by LC-MS between pup and adult female urine. (C) Enrichment of compounds in adult and pup urine that were the top 12 significantly enriched compounds in pup urine, as measured by LC-MS. (D) Supervised clustering of the all compounds analyzed by GC-MS between pup and adult female urine. (E) Enrichment of compounds in adult and pup urine that were the top 12 significantly enriched compounds in pup urine, as measured by GC-MS.

Further, to test whether the observed preference extends to non-social odors, virgin females and PPD5 mothers were presented with a nonsocial odor, acetophenone. Neither group exhibited a specific preference for this odor in time spent, latency to first interaction or cotton movement (Figure 1N–O, Figure S1C). In fact, mothers showed a significant reduction in the number of interactions with acetophenone (Figure S1C2). These results suggest that the shift in odor preference observed in PPD5 mothers is specific to pup urine and does not reflect a generalized change in olfactory attraction.

Finally, to rule out the possibility that the observed preference for pup urine was driven by the wet texture of the cotton rather than the odor itself, PPD5 mothers were tested with a choice between wet and dry cotton. The results did not show a significant preference for wet cotton, confirming that the attraction to pup urine is mediated by olfactory cues rather than the cotton’s tactile properties. Additionally, average nose speeds did not differ significantly between virgin females and mothers across all conditions (Figure S1D). Taken together, these findings demonstrate that postpartum mothers, but not virgin females, develop a specific olfactory preference for pup urine, reflecting a maternal-specific shift in pup odor preference.

### Pup urine is distinct from adult urine

What makes pup urine special? To explore how it differs from adult urine and what might underlie the observed maternal preference, we analyzed its chemical composition. We used gas chromatography-mass spectrometry (GC-MS) to identify volatile organic compounds and liquid chromatography-mass spectrometry (LC-MS) to profile untargeted metabolomics (Figure 2A).

For LC-MS, urine samples were collected from 5-day-old male and female pups and pooled into three technical replicates, as well as 10-week-old virgin females in diestrus and proestrus stages, two estrus cycles characterized with low and high estrogen levels, respectively. The metabolic profile of pup urine was distinct from that of adult virgin females (Figure 2B). Differential analysis of pup and adult female urine revealed 140 compounds with significantly more enrichment in pup urine, and 208 compounds significantly more enriched in adult urine (Table S2). Of the compounds with significantly more enrichment in pups, Proline is the most abundant compound enriched in pup urine (Figure 2C).

For GC-MS, urine samples were collected from 5-day-old male pups, 5-day-old female pups, 10-day-old male pups, 10-day-old female pups, mothers, adult virgin females, and adult males. Unsupervised clustering reveals high similarity of all identified volatile compounds in pup urine, regardless of sex or stage (Figure 2D). Of the 67 identified compounds, 25 are significantly more enriched in pup urine, compared to all adult female samples (Table S3). Of these 25 compounds enriched in pups, those with the highest fold change in enrichment include HMH, 5-methyl-2-hexanone, 4-propyl-2,3-dihydrofuran, 3,4-dimethyl-2-hexanone, 2-acetyl-1-pyrroline, 1-decanol, an unknown compound, substituted exo-brevicomin, possible 3-fluoro-p-anisidine, methyl methylthiomethly disulfide, hexanoic acid, and a-farnesene (Figure 2E). Taken together, these analyses revealed that pup urine has a distinct chemical signature with specific metabolites and volatile compounds enriched, some of which may function as olfactory or contact-mediated cues driving maternal preference.

### Olfaction and contact chemosensation are both important for maternal preference of pup urine

Are both volatile and non-volatile components necessary for maternal preference toward pup urine? To address this, we next investigated the roles of olfactory and contact chemosensation. To disrupt olfaction, we administered intraperitoneal injections of methimazole (MMZ), which causes rapid and reversible ablation of the main olfactory epithelium (MOE)^10^ (Figure 3A-B). MMZ-injected mothers exhibited significantly impaired pup retrieval performance compared to PBS-injected controls, confirming that this manipulation was effective and that volatile cues are necessary for guiding pup retrieval behavior (Figure 3E). In tri-chamber odor preference tests, control PBS-injected mothers showed a significant preference for pup urine–soaked cotton, as indicated by both total time spent and a preference index significantly greater than zero (Figure 3C). In contrast, MMZ-injected mothers showed no preference by either metric (Figure 3C). MMZ treatment did not significantly affect average speed compared to PBS-injected controls (Figure 3D), indicating that general locomotion and exploratory behavior remained intact. Notably, PBS-injected mothers in this batch showed a faster initial approach to the pup urine stimulus compared to earlier cohorts (compare latency to first interaction in Figure S2B to Figure 1H), suggesting rapid attraction to pup odors. MMZ-injected mothers, however, showed no such latency bias (Figure S2B). Moreover, PBS-injected mothers in this cohort showed no significant bias in total number of interactions with pup urine versus control cotton, and movement of pup urine versus control cotton compared to earlier cohorts (compare Figure S2A & S2C to Figure 1G &1I). These discrepancies likely reflect batch-to-batch variability in behavioral expression. Nevertheless, the overall pattern that mothers exhibit a preference for pup urine, remained consistent.

**Figure 3.**
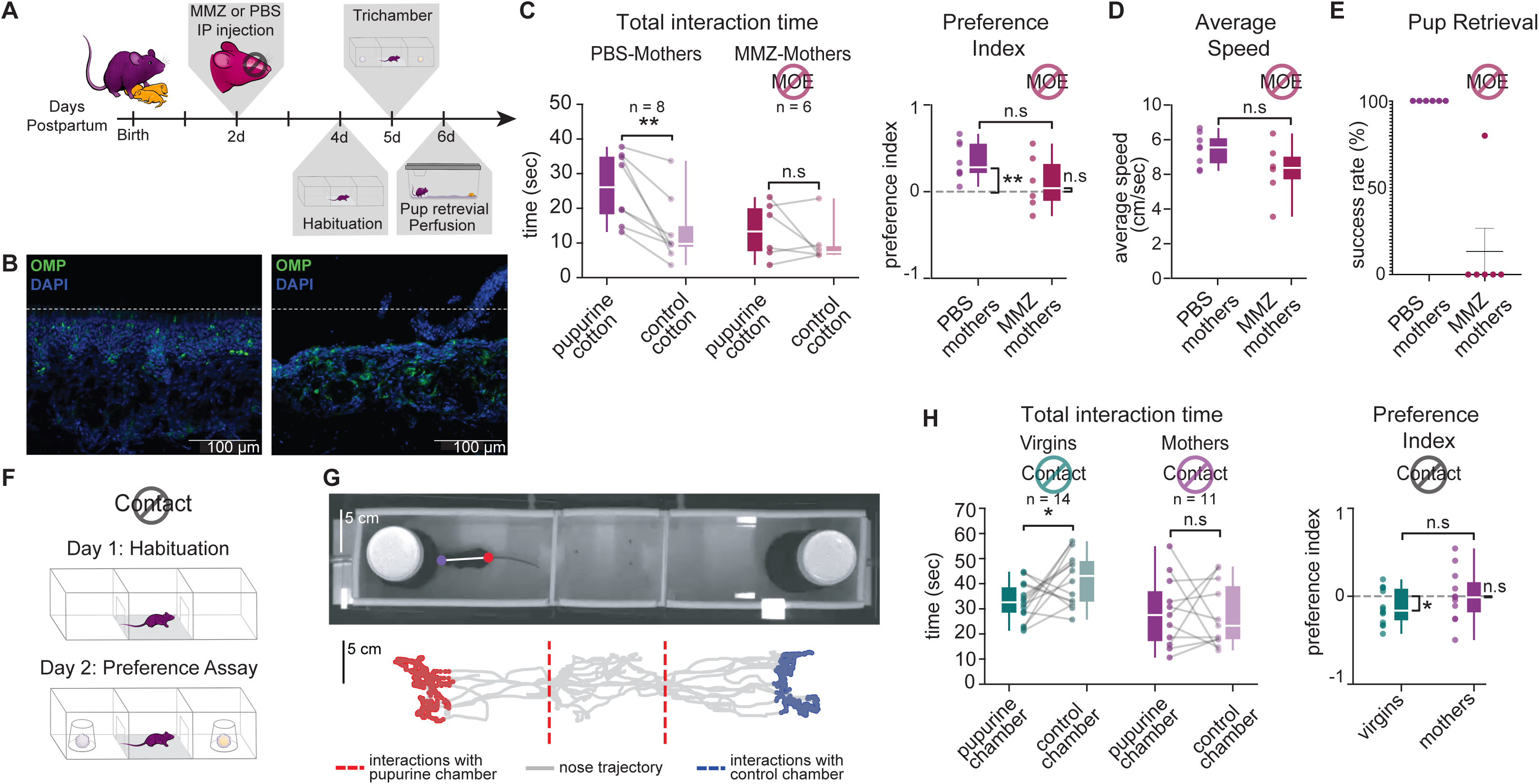
**Olfaction and contact chemosensation are both important for maternal preference of pup urine.** (A) Behavioral paradigm used to ablate the main olfactory epithelium (MOE) and assess pup urine preference and pup retrieval success rate. (B) Confocal images of the MOE focusing on the zone 1 epithelial layers of mothers injected with PBS (left) and MMZ (right). Olfactory marker protein (OMP, green) labels mature olfactory sensory neurons; DAPI (blue) marks nuclei. (C) Left: Total interaction time with pup urine-soaked versus blank cotton in PBS-(*n* = 8) and MMZ-injected (*n* = 6) PPD5 mothers. Right: Preference index calculated from the left panel (see Methods). (D) Average nose movement speeds of PBS- and MMZ-injected PPD5 mothers during the pup urine assay. Groups compared using the Mann–Whitney U test. (E) Pup retrieval success rates in PBS- and MMZ-injected mothers over 10 trials (*n* = 6 per group). (F) No-contact odor preference tri-chamber assay design, where cotton samples were enclosed in mesh cups to block direct contact. (G) Top: Example camera frame from a representative session. Bottom: Nose trajectory (gray) from a single 2-min session. Red and blue dots mark interactions with pup urine and blank cotton chambers, respectively. (H) Left: Total interaction time in pup urine versus blank cotton chambers in PPD5 mothers (*n =* 11) and virgin females (*n =* 14). Right: Corresponding preference index scores (see Methods). See Figure 1 legend for details of statistical analyses and significance conventions.

Next, to restrict contact chemosensation, we enclosed the pup urine– and control cotton– soaked samples inside wire mesh cups, preventing direct physical access while allowing volatile cues to diffuse (Figure 3F). We quantified interactions with each chamber (Figure 3G) and found that virgin females avoided the chamber containing pup urine cotton, while PPD5 mothers showed no preference between the two chambers (Figure 3H). Neither group showed significant differences in the total number of interactions or latency to first interaction to either cotton samples. These results suggest that virgin females may exhibit an aversive response to initial exposure to pup urine volatiles – a response that is absent in postpartum mothers. Notably, there was no significant difference in preference index between virgins and mothers in this condition, nor between PBS-injected and MMZ-injected mothers in the olfactory disruption condition, further highlighting the limitations of the preference index in capturing subtle or context-dependent behavioral differences.

### Development of pup urine preference in PPD5 mothers requires both hormonal and chemosensory cues

To determine whether pup urine preference is established before birth or develops postpartum (Figure 4A), we first tested pregnant females at E18.5. Pregnant females did not exhibit a preference for pup urine in the tri-chamber assay (Figure 4B). PPD30 mothers were also tested in the same paradigm to assess whether pup urine preference observed at PPD5 persists once the pups are older and are no longer in the nest. PPD30 mothers did not show a preference for pup urine (Figure 4B). Furthermore, this preference appears to be abolished by PPD30 after the pups have left the nest. These findings suggest that the preference for pup urine develops between late pregnancy and early postpartum, a critical window encompassing parturition, pup exposure, lactation, and the onset of maternal behaviors. To assess whether hormone-guided changes, in the absence of pup exposure, are sufficient to induce pup urine preference, we tested PPD5 mothers whose pups were removed on PPD0. PPD5 mothers not co-housed with their pups (“pup-deprived”) did not show a specific preference for pup urine (Figure 4C). This supports the hypothesis that pup exposure is necessary for the development of pup urine preference.

**Figure 4.**
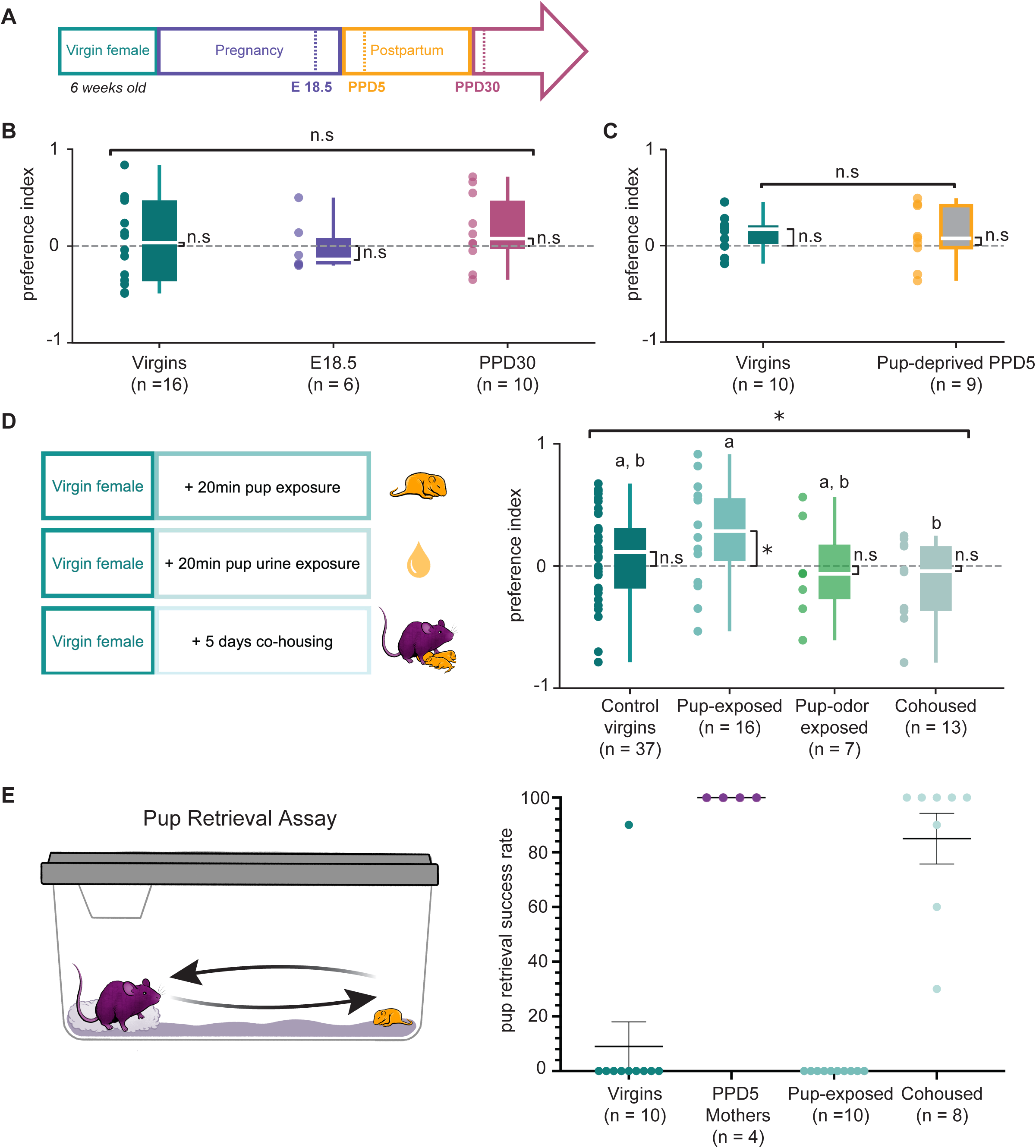
**Development of pup urine preference relies both on internal state and experience.** (A) Timeline of key stages of motherhood. (B) Preference index for pup urine vs. blank cotton in virgin females (n = 16), pregnant females at embryonic day 18.5 (n = 6), and postpartum day 30 mothers (PPD30; n = 10). (C) Preference index for pup urine vs. blank cotton in virgin females (n = 10) and PPD5 mothers with no pup exposure (Pup-deprived PPD5; n = 9). (D) Preference index for pup urine in virgin females naïve to pups (Control virgins; n = 37), virgin females exposed to pups for 20 minutes prior to testing (Pup-exposed; n = 16), virgin females exposed to pup odor for 20 minutes prior to testing (Pup odor-exposed; n = 7), and virgin females co-housed with a mother and her litter from postpartum day 0 to day 5 (Co-housed; n = 13). Statistical significance was assessed using a Kruskal–Wallis test followed by Dunn’s post hoc test. Results of statistical analyses are represented as letter codes such that groups that are not statistically significant from each other have the same letter code. (E) Pup retrieval test results for virgin females (n = 10), PPD5 mothers (n = 4), pup-exposed (n = 10), and co-housed virgin females (n = 8). See Figure 1 legend for details of statistical analyses and significance conventions.

Next, to investigate whether pup exposure alone can induce a preference for pup urine, we exposed virgin females to pups prior to testing. Previous studies have shown that virgin females co-housed with an experienced mother and her litter can develop alloparental behaviors such as pup retrieval.^11–14^ Interestingly, virgin females exposed to pups for an extended duration (five days with the mother and litter) did not exhibit a significant preference for pup urine (Figure 4A). In contrast, those exposed to pups for a short duration showed a significant preference relative to chance (i.e., a preference index significantly different from 0 and greater total interaction time with pup urine–soaked cotton), as well as a significantly higher preference index compared to the extended exposure group (Figure 4A). This pattern may reflect a novelty-driven response following a second, brief exposure to pup-related olfactory cues. Notably, virgin females exposed only to pup urine (without pups) for 20 minutes did not show a preference (Figure 4A). Taken together, these findings suggest that pup urine preference is not solely driven by pup exposure or familiarity, but rather requires the physiological changes associated with gestation and parturition. Overall, these results suggest that the development of pup urine preference does not arise solely from long familiarity with pups or pup odor but rather requires the physiological changes associated with gestation and parturition.

Interestingly, virgin females co-housed with a mother and her pups demonstrated robust pup retrieval behavior despite lacking a preference for pup urine, indicating pup urine preference is not correlated with successful pup retrieval (Figure 4E). It remains unclear whether these virgin females engaged in other maternal behaviors, such as licking the anogenital region, which could maybe contribute to the development of pup urine preference.^9^ Overall, this finding suggests that pup retrieval and urine preference are mediated by distinct mechanisms, with retrieval potentially driven by auditory or tactile cues rather than olfactory inputs.

## Discussion

Parental behavior is often described as the output of evolutionarily sculpted neural circuits - hardwired and activated by a switch in physiological state. While this framework has helped define key brain regions and hormonal correlates of parenting, our findings suggest that maternal responses to pup-associated cues are not purely innate once hormonal priming occurs. We show that neither hormonal priming alone (as in pup-deprived mothers) nor pup experience alone (as in co-housed virgins) is sufficient to elicit a preference for pup urine. Instead, this preference appears to emerge from a convergence of internal state and sensory experience, suggesting the existence of a gating mechanism that permits pup-related cues to be reinforced only under specific maternal conditions. Moreover, we find that this attraction is not simply a feature of motherhood per se, but highly stage-dependent PPD5 mothers, but not PPD30 females, exhibit this preference, coinciding with peak lactation and nest-building. This fine-grained specificity is often underappreciated, and challenges the assumption of a unitary maternal brain. Supporting this, we observe that pup urine preference dissociates from pup retrieval behavior, retrieval can occur in co-housed virgins without any preference for pup odor, suggesting that distinct neural pathways underlie different facets of maternal behavior. Finally, this preference is shaped by both volatile and non-volatile components of pup urine: volatiles may act as distal cues that attract mothers to the source, while non-volatiles, accessible through direct contact, may serve as reinforcers that strengthen the behavioral response.

Why might it matter that mothers exhibit a preference for pup urine at this stage? One possibility is that pup urine provides context-specific information about the presence and location of pups, enhancing maternal responsiveness in a crowded nest or under threat of environmental disturbance (like infanticidal males). The increased movement of pup-urine-soaked cotton by PPD5 mothers may reflect not just olfactory attraction, but a functional drive to maintain the hygiene and olfactory distinctiveness of the nest. This is consistent with theories that maternal animals actively shape the sensory environment to optimize pup survival, for instance by removing soiled bedding that could attract predators or by reinforcing pup-associated odors to guide future maternal attention. The fact that mothers, but not virgins, override their baseline aversion to pup volatiles further suggests that motherhood entails a state-dependent loss of aversion, which allows infant stimuli to become attractive and behaviorally salient. Such flexible modulation of chemosensory cues likely enhances maternal efficiency and promotes pup survival.

Our behavioral data offers key insights into the neuroethological architecture of maternal motivation. This raises the question: how might such circuit-level plasticity modulate olfactory perception in mothers, allowing formerly neutral or aversive cues to gain motivational salience? <u>Alternatively, how might plasticity of the main olfactory epithelium during the transition to</u> <u>motherhood play a role in modulating olfactory perception?</u> The dual requirement of hormonal priming and pup experience for odor preference points to a gating mechanism, whereby pup cues are only reinforced under a set of physiological and experiential conditions. Volatile and non-volatile pathways may feed into distinct but converging circuits that integrate state and stimulus to drive behavior. This work provides a foundation for future mechanistic investigations into how the maternal brain integrates state- and experience-dependent signals to represent, update, and reinforce pup-associated chemosensory cues that guide behavior.

## Resource Availability Lead contact

Further information and requests for resources should be directed to, and will be fulfilled by, the lead contact, Bianca Jones Marlin

## Materials availability

No new materials were generated for this study.

## Data and code availability

- Code for reproducing all analyses is available at https://github.com/BJMarlinLab/Andreu-2025
- All data used in this study and any additional information required to reanalyze the data reported in this paper is available from the lead contact upon request.

## Acknowledgments

The authors thank the Biomarkers Core Laboratory at Columbia University for their assistance in generating and processing liquid chromatographic mass spectrometry data. The authors thank the Monell Institute for their assistance in generating and processing gas chromatographic mass spectrometry data. The authors thank Dr. Matteo Farinella for creating artwork used in this publication.

## Author contributions

Conceptualization: VA, NEHM, BJM Methodology: VA, NEHM, EJL, BJM

Investigation: VA, RS, NEHM, EJL, BJM Visualization: VA, RS, NEHM, EJL, DF, AS, BJM Funding acquisition: BJM

Project administration: VA, BJM Supervision: BJM

Writing – original draft: VA, BJM

Writing – review & editing: VA, RS, DF, AS, BJM

## STAR Methods

### Key resources table

| REAGENT or RESOURCE | SOURCE | IDENTIFIER |
| --- | --- | --- |
| **Antibodies** |  |  |
| anti-OMP | Abcam | ab87338 |
| **Experimental models: Organisms/strains** |  |  |
| Mouse: C57BL/6J | The Jackson Laboratory | #000664 |
| **Chemicals, peptides, and recombinant proteins** |  |  |
| Acetophenone |  |  |
| Mineral oil | Sigma-Aldrich | M5904-500ML |
| Methimazole | Sigma-Aldrich | 46429-250MG |
| Paraformaldehyde | Electron Microscopy Sciences | 15710-S |
| Phosphate Buffer Solution | Life Technologies Corporation | 10010-031 |
| Sucrose | Sigma Aldrich | S0389 |
| OCT compund | Fisher | 23-730-571 |
| Triton X-100 | Sigma-Aldrich | X100 |
| DAPI | Invitrogen | D1306 |
| Vectashield Plus mounting medium | Vector Labs | H-1900 |
| **Software and algorithms** |  |  |
| Python version 3.9.1 | Python Software Foundation | https://www.python.org |
| DeepLabCut | Mathis et al | https://github.com/DeepLabCut/DeepLabCut |
| Segment Anything | Kirillov et al | https://github.com/facebookresearch/segment-anything |
| Code for behavioral analysis | This paper | https://github.com/rs3966/rai_marlinlab |
| Numpy | Harris et al | https://numpy.org/ |
| Pandas | Mckinney et al | https://pandas.pydata.org/ |
| Scikit-learn | Pedregosa et al | https://scikit-learn.org/stable/ |
| Scipy | Virtanen et al | https://scipy.org/ |
| Statsmodels | Seabold et al | https://www.statsmodels.org/stable/index.html |
| Matplotlib | Hunter et al | https://matplotlib.org/ |
| BORIS | Friard et al | https://www.boris.unito.it |
| **Other** |  |  |
| Tri-chamber design files | Marlin Lab | bit.ly/4IJTkDt |
| Fisher Superfrost Pluss glass slides | Fisher | 12-550-15 |

### Experimental model and study participant details

#### Mice

All procedures were approved by the Columbia University Institutional Animal Care and Use Committee under protocol #AABP3561. All mice in this project were from a C57BL/6J background and were obtained from the Jackson Laboratory. All animals were housed with a 12 hr light/12 hr dark cycle and fed ad libitum.

### Method Details

#### Experimental Design

Groups of experimental mothers were formed by co-housing virgin female mice with 6-week-old males for two weeks. After this period, the males were removed, and the females were housed with her pups. Age-matched virgin females were also co-housed before testing and single-housed the day before habituation to the tri-chamber apparatus.

#### Odor preference behavioral testing

Both mothers and virgin females were introduced to a custom-built acrylic three-chamber box (the tri-chamber) when mothers reached postpartum day 4 and were tested with odors on postpartum day 5. Habituation consisted of 5 minutes in the center chamber, followed by 10 minutes of unrestricted exploration within the entire apparatus. The following day, odor preference behavior was assessed in the same setup. Mice were first accustomed to the center chamber for 2 minutes and then allowed to explore the entire apparatus for 3 minutes before testing. Test and control odors were pseudorandomized to either side of the tri-chamber and 200 μl of each pipetted into two identical, clean cotton balls. Mouse urine was collected from 5-day-old pups (male and female combined) and adult female mice, stored at −80°C, and kept on ice on the day of testing. Acetophenone was diluted to 1% v/v in mineral oil. To ensure odors remained localized, the three-chamber box was equipped with vacuum ports in the center chamber and near the doorways of the side chambers. In the no-contact version of the assay, the same tri-chamber setup was used, but the cotton samples were enclosed in mesh cups to prevent direct physical contact while allowing odor detection. All other procedures, including habituation, testing timeline, and odor preparation, remained identical.

#### Odor preference behavioral quantification

We first segmented the tri-chamber region from each behavioral video using the Segment Anything tool (https://github.com/facebookresearch/segment-anything)^15^ in an automated fashion. Masks were generated from the first or last frame of each video and selected based on area-specifically, the one whose area matched the known area of the tri-chamber in our videos. These masks were then used to crop the original videos to include only the tri-chamber apparatus. Segment Anything was run again to isolate the middle chamber within the tri-chamber apparatus, selected based on its area and central location in the video frame. This provided the x-coordinates of the left and right walls, which were used to define chamber boundaries–critical for determining when the animal exited the center and thus for identifying the start of each trial.

Nose and tail-base positions of the animal, along with the center points of the left and right cotton samples, were tracked using DeepLabCut.^16^ Sudden speed jumps (>100 pixels/frame) in any of these points were flagged as tracking errors and corrected by merging them with neighboring values. The test period was defined as beginning when both cotton samples were simultaneously detected for at least 30 seconds. A trial start was marked when the tail-base exited the middle chamber (i.e., crossed either the left or right chamber boundary) after the onset of the choice period. Cotton interaction was manually scored by blinded experimenters using BORIS (Behavioral Observation Research Interactive Software), and these observations were used to determine the spatial and angular thresholds for automated interaction detection. An interaction with a cotton sample was identified when two conditions were met: 1) the Euclidean distance between the nose and cotton center was <35 pixels (∼4 cm), 2) the angle between the vectors nose-to-tail-base and tail-base-to-cotton center was <35°. For the no-contact experiments, only the nose and tail-base of the animal were tracked. The midpoints of the left and right chambers were manually annotated. Interaction with a chamber was defined as occurring when the nose was within 85 pixels of the chamber midpoint (approximately 1.5 times the chamber radius), and the angle formed between the nose–tail-base and tail-base–chamber midpoint vectors was less than 90 degrees. (These spatial and angular thresholds were consistent with manual observations of interaction behavior in the no-contact setup, where the mesh cups spanned the full width of each chamber). In both contact and no-contact experiments, interaction bouts shorter than 10 frames were excluded, and bouts occurring within 5 frames of each other were merged into a single continuous interaction.

Mice were required to sample each cotton sample for a minimum of 100 ms to be included in the analysis. We restricted our analysis to the first 2 minutes of the test period, during which the behavioral responses were most robust. Total interaction time was defined as the cumulative duration of all valid interaction bouts during this 2-minute period. The number of interaction bouts was also quantified over this period. Preference index was calculated as (Interaction time with stimulus cotton – Interaction time with control cotton) / (Interaction time with stimulus cotton + Interaction time with control cotton). Latency to first interaction was defined as the time (in seconds) from the start of the test period to the first frame in which the animal engaged in an interaction with either stimulus or control cotton, as determined by the spatial and angular criteria described above. Nose speed was calculated as the frame-by-frame Euclidean distance (in pixels) traveled by the nose keypoint, converted to cm/s using a known pixel-to-distance conversion factor. Cotton movement distance was similarly computed by summing the frame-by-frame movement of the cotton center keypoints and converting the result to cm.

#### Pup retrieval testing

Each session consisted of 10 trials performed in the mice’s home cage. The animals were first placed in their home cage with nesting material and allowed at least 5 minutes to acclimate before testing began. A pup, aged postnatal days 4 to 6, was placed in a corner of the arena opposite the nesting material. The experimental female was given 10 trials, each lasting 2 minutes, to retrieve the displaced pup and return it to the nest. If the pup was not retrieved within 2 minutes, it was returned to the home cage with its dam, and the trial was recorded as a failure. If the pup was successfully retrieved, the time to retrieval was recorded, and the pup was then returned to its home cage. The next trial began with a new pup placed in the opposite corner.

After completing all 10 trials, the pups were returned to their home cage with their dam. The pup retrieval success rate was calculated by dividing the number of successful retrievals by the total number of trials (10).

#### GC-MS analysis

GC-MS analysis was performed by the Monell Chemical Senses Center. Urine (25 µL) was added to a 20-mL headspace vial and spiked with 10 µL of 3 µg/mL acetophenone-d5 internal standard. Vials were sealed and analyzed using gas chromatography-mass spectrometry (GC-MS) with solid-phase microextraction (SPME). Automated collection was done at 37°C for 10 minutes, with a DVB/Carbon-WR/PDMS Arrow® fiber, followed by thermal desorption at 230°C for 2 minutes. GC analysis used a Thermo Scientific ISQ MS with a Stabilwax-DA column and helium carrier gas. The oven temperature ranged from 40°C to 230°C. Mass spectrometric detection was in scan mode (33–400 m/z) with a 42-minute runtime. Data were processed using Metalign and MSClust to identify peaks and calculate peak abundance. Features were filtered by comparing peak responses from biological samples to blank QC samples, with peaks retained when the response exceeded the 95th percentile of blank samples. Normalized peak abundances were calculated by dividing by the abundance of an internal spike-in standard. A z-normalization was then applied to assess concentration of enriched compounds, and Multiple Kolmogorov-Smirnov tests were performed to evaluate differences in enrichment between groups with a FDR of 1%. Differential enrichment was calulcated using DEqMS.^17^ Heatmaps were generated using ComplexHeatmaps in R.^18^

#### LC-MS analysis

LC-MS was performed by the Biomarkers Core Laboratory at Columbia University. Untargeted Metabolomic profiling was performed using (HRAM) mass spectrometry, as previously reported.^19,20^

Metabolites were extracted from 50ul of human plasma samples using protein precipitation with acetonitrile containing a mixture of 9 stable isotope internal Standards. Ten microliters of metabolite extracts were analyzed using a high-resolution accurate-mass (HRAM) platform consisting of Vanquish DUO ™ UHPLC system equipped with dual split sampler configuration coupled to a Exploris E240 Orbitrap mass spectrometer (Thermo Fisher Scientific, San Jose, CA, USA).

Chromatographic separation was performed in triplicate by HILIC chromatography under positive ion mode and RP chromatography under negative ion mode, both at 60°C. HILIC separation was achieved on a Waters XBridge BEH Amide XP column (2.1x 50mm. 2.5 μm) and gradient elution with 0.2% formic acid in water (solvent A) and acetonitrile (solvent B). The initial 75% B at 0.35 mL/min was kept for 1.5min, decreased linearly to 20% B at 4 min with a flow rate increase to 0.4 mL/min and a final hold of 1 min. Reverse-phase separation was performed on a C18 column (Higgins Targa C18 2.1 × 50 mm, 3 μm) with 1mM ammonium acetate in water (solvent A) and acetonitrile (solvent B). The initial 35% B at 0.4 mL/min was increased linearly to 95% B at 1.5 min and held for the remaining 3.5 min with a flow rate of 0.5 mL/min. The HRMS was operated in full scan acquisition mode at 120,000 resolution and a full-scan data-dependent MS2 acquisition mode at resolutions of 60,000 (full scan) and 7500 (dd-MS2 scan) in both positive and negative polarity to acquire the spectral data. The HRMS source parameters were as follows: spray voltage 3.5 kV (+ESI), 3.00 KV (-ESI); capillary temperature 300 ℃; sheath gas flow rate 45; Aux gas 25 AU (+ESI), 15 AU (-ESI); sweep gas flow rate 1 AU; Aux gas heater temperature 250 ℃.

Raw data files acquired through the Xcalibur software (version 4.1, ThermoFisher Scientifc, MA, USA) were processed using the Compound Discoverer software (version 3.3.1, ThermoFisher Scientifc, Waltham, MA, USA). The workflow utilized an adaptive curve model with a 1-minute maximum shift, 5 ppm mass tolerance, and a 3 S/N threshold for retention time alignment. Peak detection required <5 ppm mass error for extracted ion chromatograms with a minimum peak intensity of 1e6. Feature annotation was achieved by using mzVault (internal ddMS2 database), mzCloud, and ChemSpider (HMDB, KEGG, LipidMAPS as databases).

Multivariate (PCA) and univariate (t-test) were performed using R to visualize sample clustering and identify features contributing to group separation. Metabolic pathway functional and enrichment analysis of the detected features was performed using MetaboAnalyst’s mummichog algorithm, using a list of significantly altered features (p-value < 0.05) and KEGG metabolic pathway database as the reference. Pathways with a p-value < 0.05 were considered statistically significant. Differential enrichment was calculated using DEqMS.^17^ Heatmaps were generated using ComplexHeatmaps in R.^18^

#### MOE ablation and histological verification

A single intraperitoneal injection of methimazole (100 mg/kg) was performed in mothers on postpartum day 2 to induce a detachment of the entire OE by postpartum day 5 (Håglin et al., 2021). Methimazole was diluted at a concentration of 20mg/mL in 1XM PBS, 10% DMSO. Control mothers were injected intraperitoneally with a vehicle control (1X PBS) at the same time point. Because of methimazole toxicity, mice were expected to lose around 10% of their total weight 1 day after injection.

#### Tissue processing

Mice were transcardially perfused with ice-cold 4% paraformaldehyde (PFA; Electron Microscopy Sciences, 15710-S) in 1X PBS. The main olfactory epithelia were surgically dissected, incubated overnight in 4% PFA, and cryoprotected in 30% sucrose (Sigma-Aldrich, S0389). After cryoprotection, the epithelia were embedded in OCT compound (Fisher, 23-730-571) and stored at −20°C until they were ready for sectioning. Tissue was sectioned into 20 µm slices, mounted directly onto Fisher Superfrost Plus glass slides (Fisher, 12-550-15), and stored at −80°C until staining. For staining, slides were acclimated to room temperature, washed three times for 5 minutes each in PBST (0.1% Triton X-100 in 1X PBS; Sigma-Aldrich, X100), and blocked with 5% Normal Donkey Serum in PBST for 30 minutes. Sections were then incubated overnight at 4°C with an antibody cocktail. The following day, slides were washed again with PBST (3 x 5 minutes) and incubated with Donkey anti-rabbit 488 secondary antibodies (Invitrogen, A21206) for 1 hour at room temperature. Afterward, slides were washed in PBST, followed by a 5-minute incubation in 1:10,000 diluted DAPI (Invitrogen, D1306) in PBST and PBS. Finally, the slides were coverslipped using Vectashield Plus mounting medium (Vector Labs, H-1900) and sealed with nail polish.

#### Confocal image acquisition

Imaging was performed with support from the Zuckerman Institute’s Cellular Imaging platform. Slides were imaged using a Zeiss Upright LSM 880 Confocal microscope and Zen Black software (Zeiss). All co-labeling images were acquired in z-stacks to ensure accuracy in co-labeling determination.

## Quantification and statistical analysis

All behavioral analyses were performed in Python. Cotton preferences within each session were compared using Wilcoxon signed-rank tests. For preference index values, comparisons against zero were made using the Wilcoxon signed-rank test, and comparisons between groups were made using the Mann–Whitney U test (for pairwise comparisons) or the Kruskal–Wallis test (for three or more groups), followed by Dunn’s post hoc test with Bonferroni correction. All other pairwise comparisons were also performed using the Wilcoxon signed-rank test, and across-cohort (unpaired) comparisons were conducted using the Mann–Whitney U test. All data did not approximate a normal distribution; therefore, non-parametric tests were applied throughout.

## Supplemental Information

- Document S1: Figures S1-S2
- Table S1: Spreadsheet with results of all statistical tests. Related to Figures 1-3 and S1-S3
- Table S2: Spreadsheet with comparisons of all compounds analyzed by LC-MS in pups and adult females.
- Table S3: Spreadsheet with comparisons of all compounds analyzed by GC-MS in pups and adult females.
- Video S1

**Video S1. Mothers spend more time inspecting and moving the pup urine-soaked cotton. Related to Figures 1A-I**.

Two-minute videos of a mother (top) and a virgin female (bottom) in the tri-chamber assay. Cotton-centers and the nose and tailbase of each mouse are tracked by DeepLabCut. Left, pup urine-soaked cotton. Right, control cotton.

---

**Figure S1.**
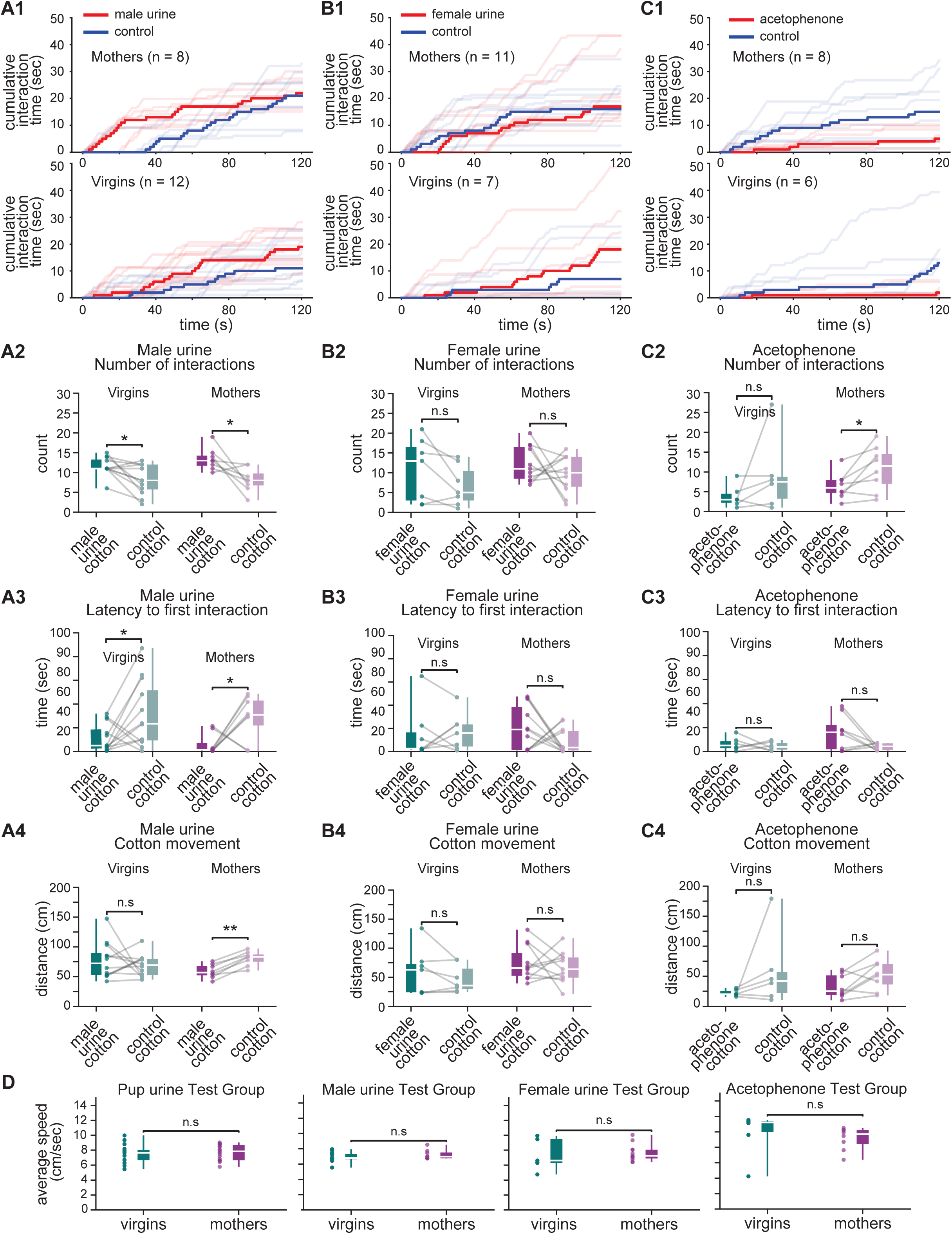
**Behavioral responses to non-pup odors in the tri-chamber assay. Related to Figure 1.** (A1) Cumulative interaction time with male urine (red) vs. blank cotton (blue) over 2 min in PPD5 mothers (top; n = 8) and virgin females (bottom; n = 12). Bold lines represent the median across animals at each frame; fine lines show individual animals. (A2-A3) Number of interactions (A2) and latency to first interaction (A3) for male urine vs. blank cotton in PPD5 mothers and virgin females. (A4) Total distance the male urine and blank cotton were moved by PPD5 mothers and virgin females. (B1) Cumulative interaction time with female urine (red) vs. blank cotton (blue) in PPD5 mothers (top; n = 11) and virgin females (bottom; n = 7). Bold lines represent the median across animals at each frame; fine lines show individual animals. (B2-B3) Number of interactions (B2) and latency to first interaction (B3) for female urine vs. blank cotton in PPD5 mothers and virgin females. (B4) Total distance the female urine and blank cotton were moved by PPD5 mothers and virgin females. (C1) Cumulative interaction time with acetophenone (red) vs. blank cotton (blue) in PPD5 mothers (top; n = 8) and virgin females (bottom; n = 6). Bold lines represent the median across animals at each frame; fine lines show individual animals. (C2-C3) Number of interactions (C2) and latency to first interaction (C3) for acetophenone vs. blank cotton in PPD5 mothers and virgin females. (C4) Total distance the acetophenone and blank cotton were moved by PPD5 mothers and virgin females. (D) Average nose movement speed of PPD5 mothers and virgin females during exposure to pup urine, male urine, female urine, and acetophenone (each vs. blank cotton). See Figure 1 legend for details of statistical analyses and significance conventions.

**Figure S2.**
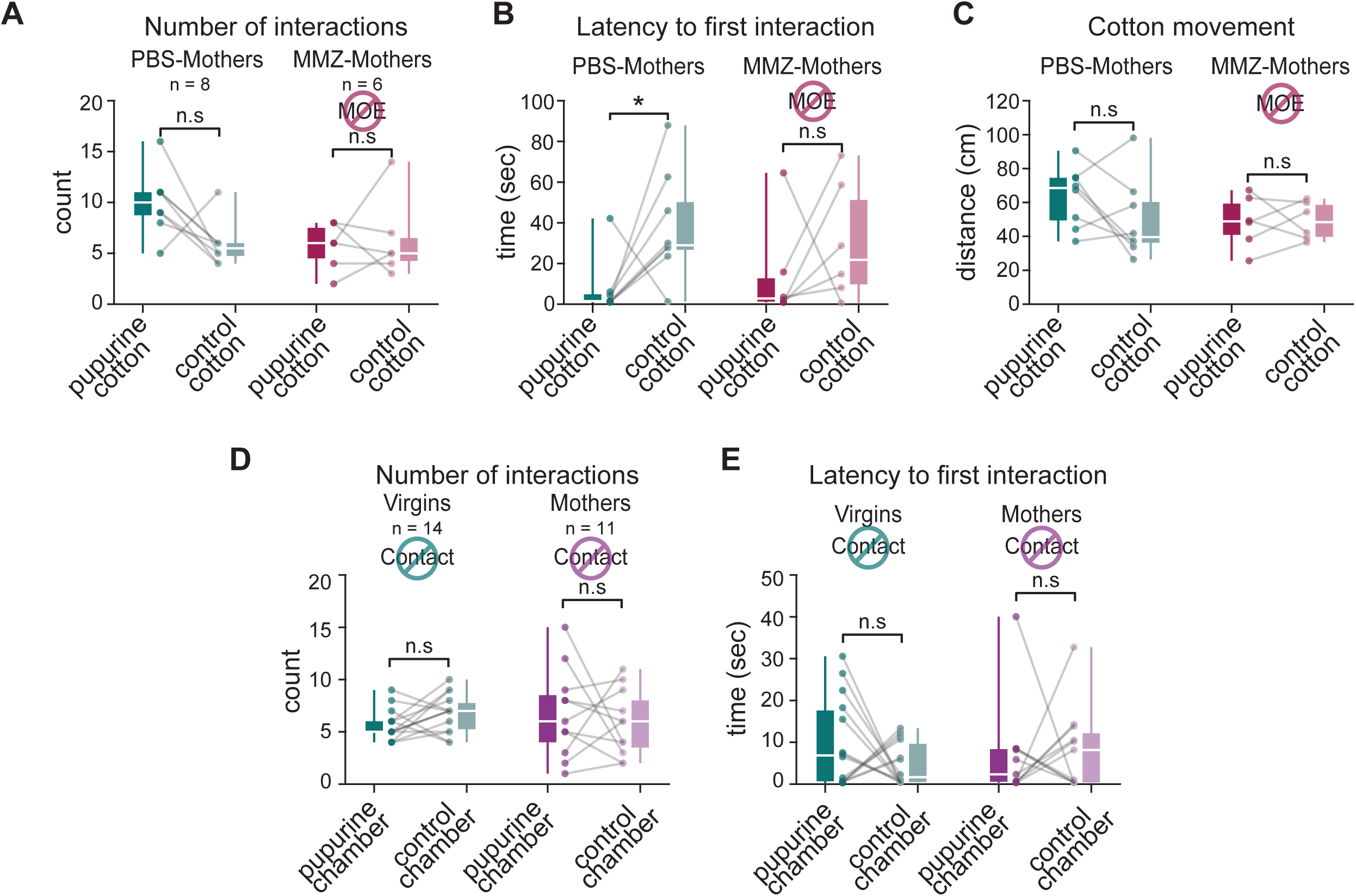
**Pup urine-directed behaviors in mothers with disrupted MOE or restricted contact to pup urine. Related to Figure 2.** (A-B) Number of interactions (A) and latency to first interaction (B) for pup urine vs. blank cotton in PBS- (n = 8) and MMZ-injected PPD5 mothers (n = 6). (C) Total distance the pup urine and blank cotton were moved by PBS- (n = 8) and MMZ-injected PPD5 mothers (n = 6). (D-E) Number of interactions (D) and latency to first interaction (E) for pup urine cotton vs. blank cotton chamber in PPD5 mothers (n = 11) and virgin females (n = 14) with no direct contact with either sample.

